# Performance characteristics allow for confinement of a CRISPR toxin-antidote gene drive designed for population suppression

**DOI:** 10.1101/2022.12.13.520356

**Authors:** Shijie Zhang, Jackson Champer

## Abstract

Gene drives alleles that can bias their own inheritance are a promising way to engineer populations for control of disease vectors, invasive species, and agricultural pests. Recent advancements in the field have yielded successful examples of powerful suppression type drives and confined modification type drives, but developing confined suppression drives has proven more difficult. This is because the necessary power for strong suppression is often incompatible with the characteristics needed for drive confinement. However, one type of CRISPR toxin-antidote drive may be strong enough and confined, the TADE (Toxin-Antidote Dominant Embryo) suppression drive. By disrupting a haplolethal target gene and a haplosufficient female fertility gene, this drive quickly eliminates wild-type alleles and eventually induces population suppression. It has been shown to perform effectively in panmictic populations. However, confinement in spatial scenarios may be substantially different. Here, we use a reaction-diffusion model to assess the performance of TADE suppression drive in continuous space. We measure the drive wave advance speed while varying several performance parameters and find that moderate fitness costs or embryo cutting (from maternally deposited nuclease) can eliminate the drive’s ability to form a wave of advance. We assess the release size required for the drive to propagate, and finally, we investigate migration corridor scenarios. Depending on the corridor size and dispersal, it is often possible for the drive to suppress one population and then persist in the corridor without invading the second population. This prevents re-invasion by wild-type, which may be a particularly desirable outcome in some scenarios. Thus, even imperfect variants of TADE suppression drive may be excellent candidates for confined population suppression.

## Introduction

Engineered gene drives are alleles that bias their own inheritance to increase transmission to offspring^1–4^. This potentially allows a gene drive to spread through a population after a modest release of drive-carrying individuals. In some cases, gene drives can be designed to modify a population, such as by spreading a cargo gene through mosquitoes to prevent transmission of diseases such as malaria. Other gene drives are designed to disrupt an essential gene as they spread, eventually resulting in population suppression or elimination. Such suppression could be used not only to combat vector-borne diseases, but also to reduce populations of invasive species or agricultural pests^1–4^.

The development of CRISPR has enabled the construction of homing suppression drives in *Anopheles* mosquitoes^5,6^ and flies^7,8^. These function by cutting the wild-type allele with Cas9 and then being copied to the cut site during homology-directed repair, thus converting germline cells to a drive homozygous state and ensuring that most offspring receive a drive allele. Because they are located in and disrupt an essential gene (usually a female fertility gene), the number of infertile individuals will increase as the drive spreads through the population, eventually resulting in population elimination if the drive is powerful enough^9,10^ and avoids formation of functional resistance alleles^10,11^. Another method for population suppression is the shred X-chromosomes, biasing the population toward males. For an X-shredder to be a gene drive, thus giving it the ability to increase in frequency in a population, it should be located on the Y chromosome (making it a Driving Y allele)^12,13^ or combined with a homing drive. However, only the latter method has been successfully demonstrated experimentally^14^ due to the difficulty of engineering genes with high expression on the Y chromosome.

Because no gene drive has yet been released, modeling is often used to assess gene drive behavior and make predictions about the outcome of a gene drive release. In most cases, stochastic models are used^15–25^, having the advantage of easily simulating random factors, complex drive mechanisms, and complex environments. However, mathematical reaction-diffusion models can potentially provide for a better fundamental understanding of gene drive properties in spatially explicit settings. Early work assessed the properties of underdominance alleles, which can also be used as modification gene drives^26–28^. Since then, such models have been used to assess the spatial spread properties of driving Y/X-shredder alleles^12^, homing drives^29^ (albeit with an embryo conversion based mechanism rather than a more accurate germline conversion mechanism), and drive recall strategies^30^. A recent study examined situations where population density changes may affect the spread of a suppression drive^31^.

Under realistic parameter ranges, homing suppression drives and driving Y systems cannot be confined to a target population. This is because these drives can substantially increase in relative frequency from very low initial frequency levels. Several designs for confined gene drive exist that have an introduction threshold, where the drive will be eliminated unless released above a particular the threshold frequency. Some of these designs have been experimentally demonstrated^32–36^, but these are generally useful only for modification rather than suppression. They can be used for suppression as a “tether” for homing suppression drives^37,38^, but this requires construction of an homing drive with high drive conversion rates, which may be difficult in many species^39–41^. One possible design, the TADE (Toxin-Antidote Dominant Embryo) suppression design can potentially overcome this issue^42,43^. Like other CRISPR toxin-antidote drives, it targets and disrupts an essential gene, in this case a haplolethal gene where two functional copies are required for viability. The drive provides a recoded rescue copy while also disrupting a haplosufficient but essential female fertility gene. This allows the drive to strongly increase in frequency and suppress the population with a power based on its total cut rate, rather than a drive conversion/homology-directed repair rate as in homing drives. It also means that the drive is frequency-dependent, giving it an introduction threshold that can potentially confine it to target populations, based on migration rates and exact drive performance parameters.

The confinement of TADE suppression drives has been studied by computational modeling in panmictic populations (includes linked 2-deme systems)^42,43^. It’s ability to suppress populations in continuous space was also studied using individual-based simulations, with a focus on its ability to avoid the “chasing” phenomenon^44^. The results of this study together with previous results on confined drives^45^ indicated that TADE suppression drive may be substantially more confined than expected in continuous space models compared to networks of linked, discrete populations. In particular, the suppressive effect of the drive may make it more difficult to spread spatially due to its frequency-dependent nature, potentially exaggerating this effect seen in previous reaction-diffusion mathematical models of suppression drives^31^.

To better understand the performance characteristics of TADE suppression drives in spatial models, we generated a reaction-diffusion framework for assessing this drive, focusing on its ability to form a wave of advance into a wild-type population. We find that less efficient drives, particularly with moderate rates of Cas9 cleavage in the embryo from maternal deposition, become substantially more confined, losing their ability to spread into a wild-type population in space. Nevertheless, TADE suppression drive is still capable of spreading in spatial populations and can often be restricted to a target population, causing successful suppression and preventing reinvasion through a migration corridor.

## Methods

### TADE Suppression drive

The TADE drive operates by targeting a haplolethal gene (usually with CRISPR/Cas9), where two copies are required for viability^42,43^. Ideally, this occurs only in the germline, converting wild-type target alleles into nonfunctional “disrupted” alleles (Figure 1). However, maternal deposition of Cas9 and gRNAs can result in additional cleavage in early embryos, which can remove drive alleles and decrease the performance of the drive. The drive allele contains a rescue version of the target gene that is recoded to prevent drive cleavage. This means that over time, wild-type alleles are removed faster than drive alleles, resulting in the drive increasing in frequency in the population. In distant-site TADE drive, the variant that we model in this study, the drive is not located at the site of its target gene. Instead, the drive is specifically located inside a haplosufficient but essential female fertility gene, which disrupts this gene, so drive homozygous females are sterile (Figure 1). With high germline cleavage rates, this allows the drive to increase in frequency to a high level, eventually resulting in large numbers of sterile females and thus a high genetic load, or suppressive power^42,43^. A same-site TADE suppression drive can be located inside its haplolethal target gene and relies on disrupting its female fertility target by use of additional gRNAs^42,43^.

**Figure 1.**
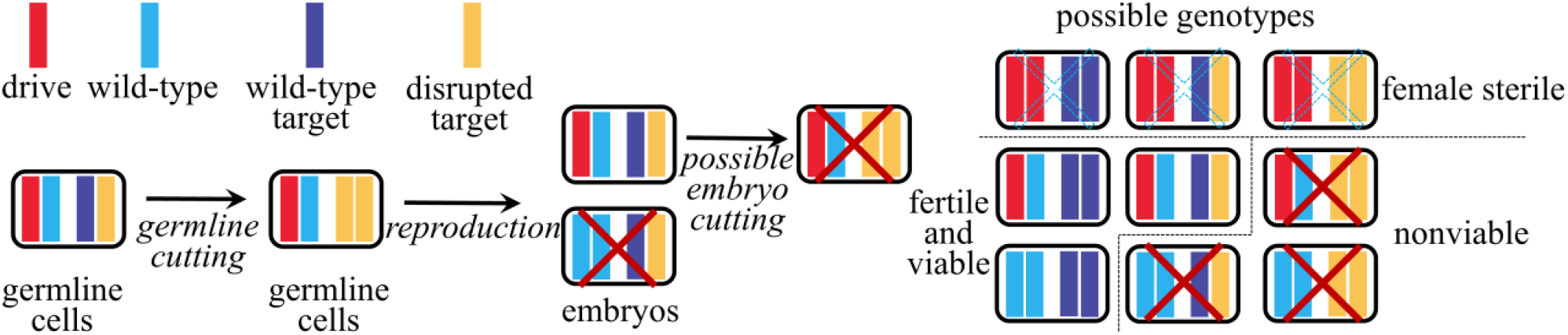
Distant-site TADE suppression mechanism and genotypes. The drive allele will cleave wild-type target alleles in the germline, converting them to disrupted alleles. Wild-type alleles in early embryos (regardless of their source) can also be converted to disrupted alleles by maternally deposited Cas9 and gRNA. If the total number of functional target gene copies (which can be drive allele or wild-type target alleles) is less than two, offspring will be nonviable (leaving six viable genotypes that are modeled directly in our simulations). Additionally, in distant-site TADE suppression, drive homozygote females will be sterile.

### Reaction-diffusion model

We simulate a population of individuals modeled by variables for each possible genotype. The whole population is initially composed of wild-type individuals (i.e. N = *X*_1_). We assume that a time interval of Δ*t* = 1 is one generation of the species modeled. We assume that when the population N is very small (i.e. N → 0), the relative growth rate of the species is β, the low-density growth rate. We set λ to be the fecundity of the species (the number of female eggs laid per generation, and we assume a constant sex ratio of 1) and μ to be the density-dependent mortality rate of the species. Thus, we have the population equation to be:

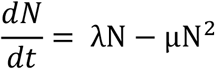

We assume the carrying capacity of the environment is 1 (making N a relative population size with respect to carrying capacity). Thus, we have λ = μ. When 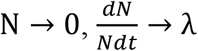. Thus, we have λ = μ = β.

To better evaluate the more realistic performance of the population behavior in a landscape, we add spatial diffusion term *D*Δ*N* to:

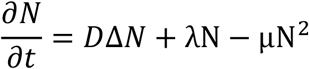

Here, Δ= 𝛻^2^ represents the Laplace operator. In this study, we mainly study 1-dimensional diffusion 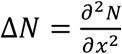, 2-Dimension diffusion 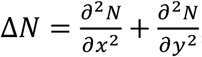, and rotationally symmetric 2-dimensional diffusion 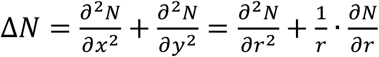.

### Drive model

We suppose for a certain genotype, females have the same population as males (i.e. *X*_*i,Female*_ = *X*_*i,Male*_ = *X*_*i*_/2). Since we suppose that each female lays λ female eggs per generation, we will have 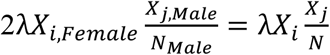 offspring if females of genotype *i* mate with males of genotype *j*. Thus, based on the TADE suppression drive strategy, we can derive our equations as follows:

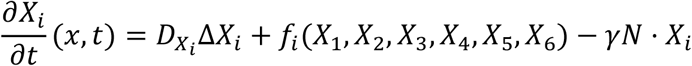

where *X*_1_(*x, t*), *X*_2_(*x, t*), *X*_3_(*x, t*), *X*_4_(*x, t*), *X*_5_(*x, t*), *X*_6_(*x, t*) are the population densities of corresponding genotype at time *t*. The total population density is N(x, t) = *X*_1_(*x, t*) + *X*_2_(*x, t*) + *X*_3_(*x, t*) + *X*_4_(*x, t*) + *X*_5_(*x, t*) + *X*_6_(*x, t*). f_i_(X_1_, X_2_, X_3_, X_4_, X_5_, X_6_) is the growth function of genotype X_i_(x, t), which is derived from the mating strategy. The specific function of f_i_(X_1_, X_2_, X_3_, X_4_, X_5_, X_6_) is in the Supplementary Appendix. We calculated the drive allele frequency as 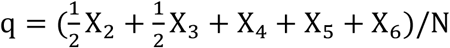. For ideal drive, we could simplify it as 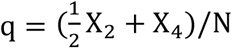. Default drive performance and ecological parameters are specified in Table 1.

**Table 1:** Parameters of the model

| Variable | Name | Default Value |
| --- | --- | --- |
| s | Germline cut rate | 1 |
| ss | Embryo cut rate | 0 |
| f | Drive allele fitness | 1 |
| $N_0$ | Normal population density | 1 |
| D | Dispersion | 0.1 |
| $\beta$ | Low-density growth rate | 5 |

### Simulation method

In this project, we use the finite difference method to simulate our reaction-diffusion model. All of our numerical simulations are performed in MATLAB R2021a. The space is a bounded domain, and we initialize the simulation using the equations below. The whole arena is initially wild-type, and we release drive heterozygotes at frequency *h* into the arena by substituting the wild-type allele with the drive allele *X*_2_ in the middle of the arena. Thus, for the 1-Dimension diffusion model, the initial condition at t=0 is:

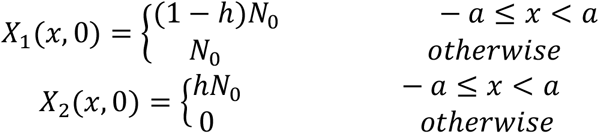

In these equations, we set the release region to have *a* = 5. For the rotationally symmetric 2-dimension diffusion model, the initial condition at t=0 is:

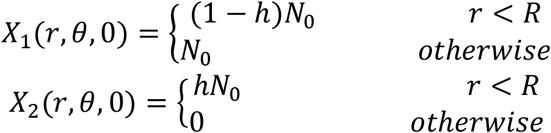

In this model, *r* varies in simulation to study how the releasing radius would affect the drive’s ability to establish (Figure 4).

### Barrier model

In the 2-Dimensional diffusion model, we consider a special type of scenario to further study the TADE suppression drive. Here, we set the arena to be two rectangular areas of length 4 and width 10 that are connected by a narrow migration corridor. We set W to be the width of the corridor and 2 to be the length of the corridor (W varies in our simulations). We set the initial condition at t=0 as follows: drive heterozygotes *X*_2_ are released at 50% frequency at the left half of the left rectangular area, and the rest of the whole arena (the right half of the left rectangular area, the right rectangular area, and the corridor area) is full of wild-type individuals.

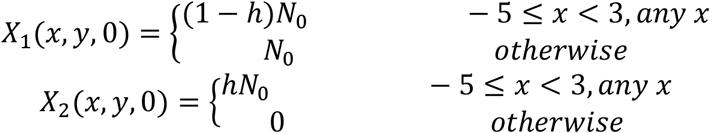

### Wave speed calculation

To calculate the speed of an advancing drive wave, we set a threshold drive allele frequency of 20% to indicate when the drive wave arrives a certain point. We set two points and recorded the time needed after initialization for the drive allele to reach each point. Then, we calculated the wave speed by:

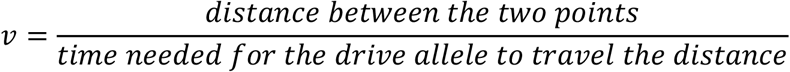

### Software

We used MATLAB to analyze data and draw figures. All MATLAB models are available on GitHub (https://github.com/jchamper/ChamperLab/tree/main/TADE-Suppression-Reaction-Diffusion).

## Results

### Advancing drive waves

Using our reaction diffusion model, we assessed the performance of TADE suppression drive waves of advance. TADE suppression drive targets a haplolethal gene where two copies are required for viability (Figure 1). These can be wild-type copies or drive alleles. The drive sites inside a female fertility gene but does not target this gene, so drive homozygote females are sterile. The drive will be able to impose a high genetic load at equilibrium if the germline cut rate is high^42,43^, meaning that it will have the power to strongly suppress the target population.

In general, if released in a limited area at sufficient quantity and with sufficiently high performance, TADE suppression drive will reach high frequency in its release area and then expand outward. For an ideal drive in a one-dimensional setting, we found that the wave quickly formed (Figure 2A). We can then clearly observe a traveling wave of the drive allele that slowly advances against the wild-type allele.

**Figure 2:**
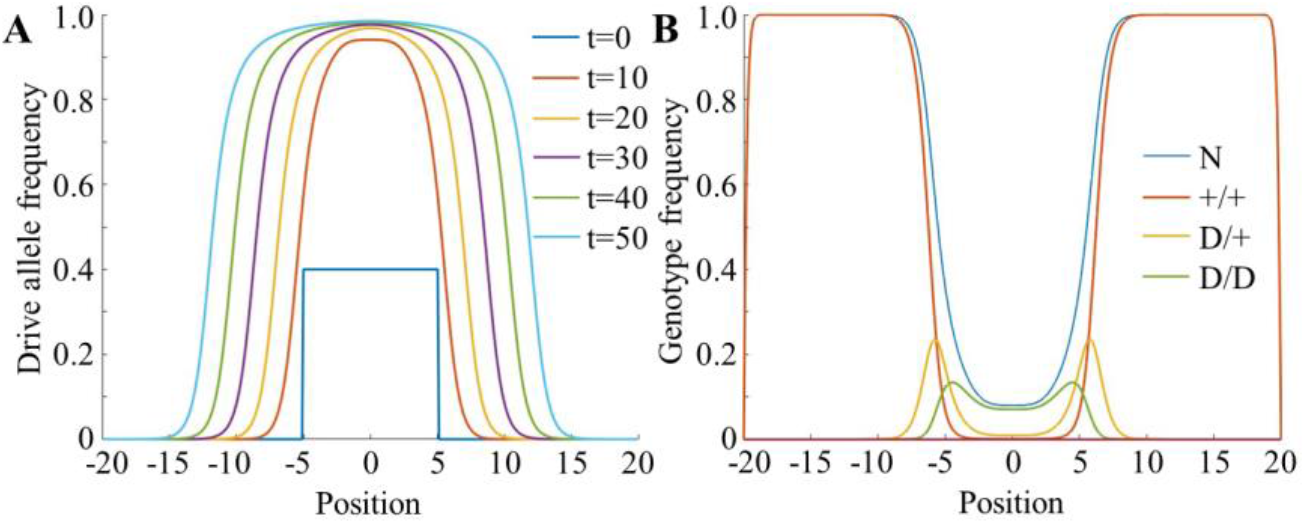
TADE Suppression drive wave shape. Drive heterozygotes are released at 80% frequency in a central area. (**A**) The wave quickly takes shape and moves outward, holding its shape as it advances. (**B**) A snapshot at t = 10 for an ideal drive. The only genotypes are wild-type (+/+), drive heterozygote (D/+, with one cut target alleles), and drive homozygote (D/D, with two cut target alleles). The population is heavily suppressed when drive homozygotes are common and eventually eliminated when drive performance is ideal.

Examining the wave in more detail, we see that an ideal TADE suppression drive has three of the six viable genotypes (Figure 2B), with other genotypes only forming when the Cas9 cleavage rates are imperfect. In the center of the drive region, the total population has been heavily suppressed. It is continually declining toward elimination (which, strictly speaking, cannot occur in our mathematical model). Drive homozygotes are present at high frequency in this area. At the leading edge of the wave are drive heterozygotes. Though they can represent a high proportion of the total number of individuals in the advancing the drive, the suppressive effect of the drive means that their individual density will not be high compared to wild-type regions. This slows the wave advance, particularly because TADE suppression is frequency-dependent, requiring high numbers of drive individuals to remove wild-type alleles.

### TADE suppression drive wave speed

The speed of the advancing drive wave is a critical parameter for assessing drive performance. It will show how rapidly the drive can spread in spatial models and is thus a fundamental property of the drive. If the drive has a speed of zero, it will not be able to advance against wild-type and will be confined to its release area. If the drive has a negative speed, it will not be able to form a wave of advance and will eventually be lost from the population (though a widespread drive release can often still allow success in spatial models in these conditions^44^).

We first investigated the effect of germline cut rate and drive fitness on the wave speed (Figure 3A). In general, fitness had a large effect. If the fitness of drives alleles was below 92.5% that of wild-type alleles, the drive would be unable to advance even if other parameters were ideal. While this represents only a modest fitness cost, note that it is a per-allele (multiplicative) fitness cost for both heterozygotes and homozygotes. A TADE suppression drive would not have a costly cargo gene, and the fitness of drive alleles without cargo would usually be close to wild-type^46^. Rescue efficiency of the haplolethal target could also likely be highly effective, preventing significant fitness costs^47^. One possible exception that could cause trouble for TADE suppression drive is somatic Cas9 expression, which would target the wild-type haplolethal gene without additional rescue, though these effects would only be significant in drive heterozygotes. Reducing the germline cut rate did not affect wave speed until it fell below about 0.75, despite the large effect of this parameter on the genetic load of the drive and its rate of increase in panmictic populations^42,43^. This is likely because lower germline cut rates reduced removal of drive heterozygotes in matings between drive individuals, allowing the density of drive individuals to be higher, which is helpful for the wave to advance.

**Figure 3:**
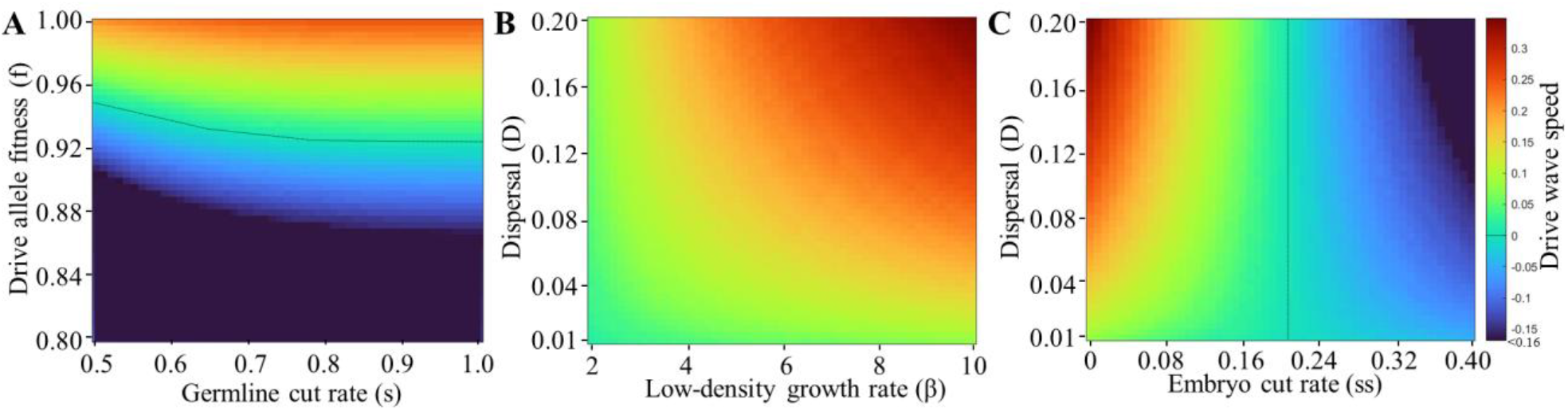
Drive wave advance speed. Simulations in one-dimensional space were initialized by releasing the drive at a sufficiently high frequency to allow it to establish. The drive wave advance speed in dimensionless units per generation of time is shown for varying (**A**) germline cut rate and fitness, (**B**) low density growth rate and dispersion, and (**C**) embryo cut rate and dispersion. Negative wave advance speeds mean that the drive will recede in favor of wild-type.

We next investigated the effect of varying the dispersal value and the low-density growth rate (Figure 3B). The dispersal, as expected, directly increased the wave speed by increasing the level of mixing between drive and wild-type, making the waves wider. Increasing low-density growth rate also directly increased the wave speed. This is accounted for by slower population suppression effect and thus higher densities of drive individuals in the whole wave of advance, an important feature for a frequency-dependent suppression drive. This result based on linear density-dependence is in contrast to other results using a nonlinear density-dependent growth curve, where reproduction only modestly increased until population density was very low, at which point the increase became more substantial^48^. This underscores the important of density-dependence for TADE suppression drive performance.

One critical parameter for TADE suppression drive is the embryo cut rate, which can increase the panmictic population introduction threshold of an otherwise ideal drive from 0% to 61% as the embryo cut rate changes from 0-100%. When varying this parameter, we found that the wave speed steadily declined as the embryo cut rate increased, an effect that was amplified by higher dispersal (Figure 3C). When the embryo cut rate was 21%, the drive wave advance speed fell to zero. Thus, if a TADE suppression drive is released in only a portion of a connected population, the embryo cut rate should be below 20% for the drive to spread on its own.

### Drive release conditions

Like other confined drives^45,49^, a TADE suppression drive must be released over a large enough area in order to successfully persist and eventually expand. To investigate this, we released TADE suppression drive heterozygotes at 80% frequency over a circular area with variable radius (Figure 4A), representing an initial deployment of the drive into an already existing population of wild-type (though this is the simplest release pattern, it is not necessarily the optimal form for a drive^49^). We measured average drive wave speed for a short interval after allowing the population to stabilize after initial diffusion of the drive allele. Thus, the wave speed is representative of the long-term behavior of the drive. We found that for an ideal drive, the critical release size is small (Figure 4B). If the drive was able to establish, then it could advance quickly. However, as the embryo cut rate increased, larger release radii were required for successful persistence of the drive, particularly at embryo cut rates of over 15%.

**Figure 4:**
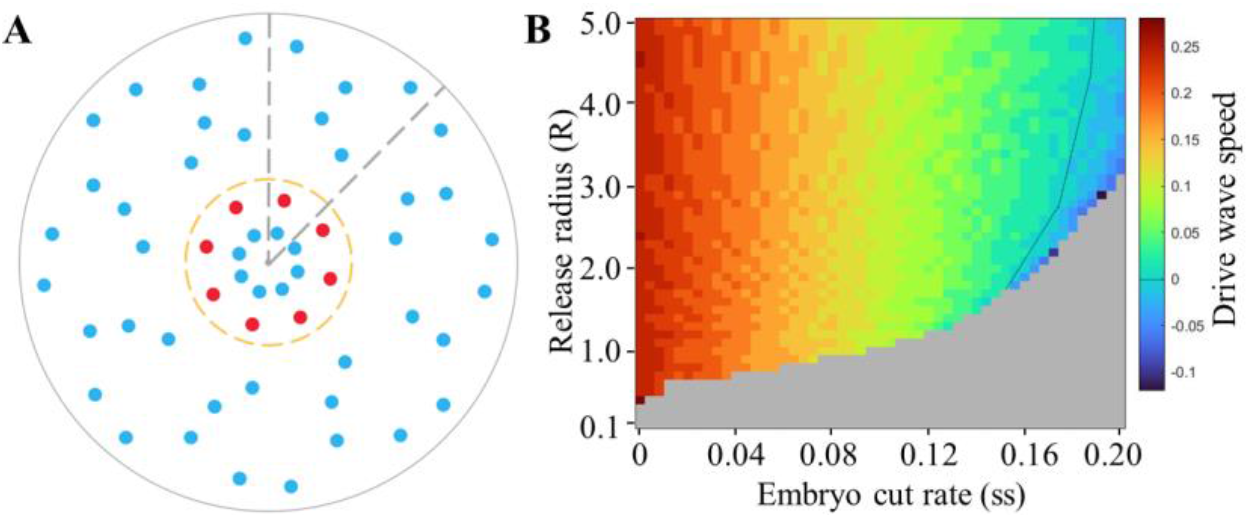
Required release characteristics for drive establishment. (**A**) Simulations in two-dimensional space were initialized by releasing drive heterozygotes at 80% frequency in a circle of variable radius. (**B**) The average drive wave advance speed in dimensionless units per generation of time is shown between generations 45 and 50 for varying release radius and embryo cut rate. Negative wave advance speeds mean that the drive will recede in favor of wild-type. The grey area shows a region of parameter space where the drive is quickly eliminated.

### Confinement of the drive in migration corridors

Drives that are able to form a wave of advance under neutral, open-field conditions in continuous space models are not necessarily always able to invade any directly connected populations^45^. This can potentially be important if there are two connected populations, but only one should be targeted by the drive. To investigate this, we simulated a migration corridor with a release of drive individuals across half of one of the two populations. We then calculated the time needed for the drive to reach the middle of the corridor (yellow dot in Figure 5A) and the time to reach the middle of the second population (purple dot in Figure 5A). The drive would always quickly spread and suppress the whole population where it was released. However, from Figure 5B-C, we can see that the drive often failed to reach high frequency in the corridor and then go on to invade the second population.

**Figure 5:**
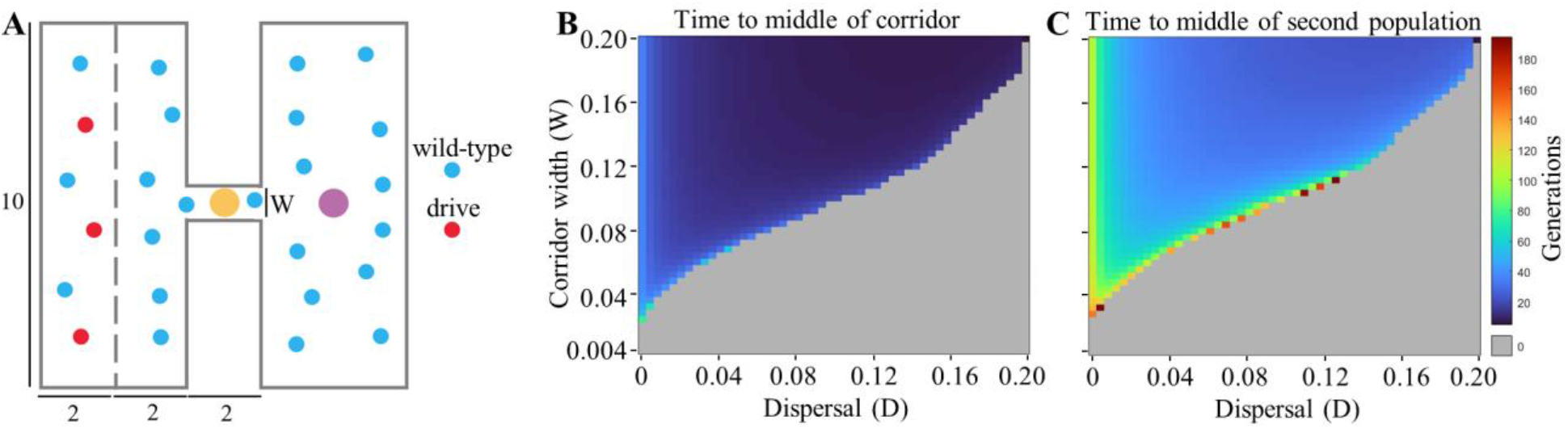
Confinement of the drive in a barrier model. (**A**) A schematic diagram of the initial state of the model. Drive heterozygotes constitute 50% of individuals left half of the left rectangular area. The yellow dot is the middle of the corridor, and the purple dot is the middle of the right rectangular area. Numbers show distances (W is the corridor width). Heatmaps are displayed for the time needed for the drive to reach 70% allele frequency in (**B**) the middle of the corridor (yellow point in panel A) and (**C**) the middle of the right area (purple point in panel A). The grey area shows a region of parameter space where the drive was unable to reach the designated point.

This is because after suppressing the first population, there are only a small number of drive individuals in the corridor and no more coming in from migration. Meanwhile, wild-type individuals can move into the corridor from the much larger second population. This allows the drive wave to be stopped in the corridor, or at least substantially delayed. This usually happens in the middle of the corridor (Figure 5B), though in some borderline cases, the drive can reach the middle of the corridor but fail to invade the second population (Figure 5C). However, in all cases where the drive failed to invade the second population, the drive wave persisted in a stationary state in the corridor, thus preventing recolonization of the first population by wild-type individuals. This is a promising result for TADE suppression drive’s capacity to achieve localized suppression, even in situations with directly connected populations.

## Discussion

The TADE suppression drive is one of a small number of possible gene drive options for confined suppression, so a better fundamental understanding of its behavior in is desirable. Such drives may be required for suppression of invasive species and agricultural pests outside their native range, or local suppression of global disease vectors as part of initial testing or where sociopolitical factors demand restriction of the drive to target populations. To assess the complexity of spatial structure, varying population size, and their interaction, we generated a flexible reaction-diffusion mathematical model to analyze this drive. We found that TADE suppression drive is capable of spreading in spatial populations despite its frequency-dependent strength and reduction of population density by drive suppression in the drive wave. However, reductions in efficiency can result in the drive being unable to persist, and the shape of the spatial structure can potentially halt the spread of the drive.

Unlike Driving Y or homing suppression drives, which have no panmictic population introduction threshold across most of their parameter space, a TADE suppression drive (which has a zero-introduction threshold in idealized form) will gain an introduction threshold with even a small fitness cost or level of embryo cutting (from maternally deposited Cas9 and gRNA) or a small fitness cost. This is because TADE suppression drive is frequency dependent. Higher drive frequencies will improve the relative increase of the drive per unit time, unlike Driving Y or homing drives, where this actually slightly decreases with increasing the drive frequency^48^. Thus, TADE suppression drive, together with most other CRISPR toxin-antidote systems, will generally have the characteristics of a pushed wave. Such waves are propagated by frequency increases in the middle of the wave where the drive is at high frequency, followed by dispersal of these new drive individuals toward the leading edge of the wave. However, if population density is lower at the middle of the wave than in the wild-type region, advancement of the drive will be inhibited^31,50^. This is a major issue that confined suppression drives have, but TADE suppression drive appears to still be able to advance if the drive has suitable parameters. However, the drive’s wave advance speed is sensitive to variation in the low-density growth rate (and the shape of the density-dependent growth curve), in contrast to other drives. Thus, higher low-density growth rates can result in substantially more efficient spread of a TADE suppression drive, making this a key parameter for TADE suppression drive, at least for some forms of density-dependence. Of course, like with other drives, if the germline efficiency is too low, the drive may lack the genetic load needed to overcome a high low-density growth rate and suppress a population^42,43^. The embryo cut rate of TADE suppression drive is also a critical parameter. Even an ideal drive will not be able to advance in continuous space models if the embryo cut rate is over 20%, which is a significant difference from panmictic models. In these simpler models, such a drive with 20% maternal embryo cutting would be only moderately confined with an introduction threshold of 27%^43^, a level where a modification underdominance drive would remain quite capable of spreading in spatial populations^45^.

Like other confined drives^28,45,51^, the release pattern is substantially more important for TADE suppression drive than unconfined systems. Even if the local introduction frequency in a central release pattern is above the panmictic introduction threshold, the drive may fail due to the geometrical disadvantage of being surrounded by wild-type (thus resulting in more inward migration of wild-type individuals than outward migration of drive individuals). Confined drives thus have more stringent initial release requirements to allow the drive to form a wave and advance outward.

While models of panmictic populations or networks of linked demes is perhaps representative of some real-world populations, other populations are likely better represented by individuals spread over continuous space. In the simple case of an even field, a drive can be considered unconfined if it is able to form a wave of advance at all. In such a case, a confined drive would always have a high introduction threshold and require releases across nearly the whole population if it is to spread^45^. However, many real populations likely have a more intricate structure, which could allow drives to be confined even if they can advance in an open field. As an example, we showed that TADE suppression drive can invade one population and induce successful suppression, but then fail to invade a second population connected by a sufficiently thin migration corridor. This is a particularly desirable result when confinement is desired. By persisting in the corridor, the TADE suppression drive avoids being eliminated, which would allow wild-type individuals in the second population to recolonize the first population. Other spatial structures such as movement barriers or density gradients could perhaps yield similar confinement results.

Engineering TADE suppression drive may prove challenging, but previous success in generating CRISPR toxin-antidote drives^32,33^ and creating rescue when targeting haplolethal genes^47^ is encouraging for their future prospects. Yet, while our model results are promising in that the drive performs well in realistic spatial environments, our math-based method does include variation in possible outcomes, particularly levels of confinement, based on drive performance parameters and local ecology. Such ecological dynamics could be affected by individual-based random factors, species lifecycles, seasonality, and other factors, so our results are complementary to previous models that can assess stochastic outcomes such as chasing^44^. To make fully accurate outcome predictions of a TADE suppression drive release, it would be essential to understand and model a species’ population dynamics specifically in the target region. Overall, though, TADE suppression drive may be an ideal candidate for restricting strong suppression to a localized region in many scenarios.

## Acknowledgements

This study was supported by laboratory startup funds from Peking University and the NSFC Overseas Youth Fund.

## Supplementary Appendix

### Growth function

We assume that the number of males and that of the females are equal for a specific species, which means males and females of species X_i_ are X_i_/2. *λ* is the fecundity of the species (the number of female eggs throughout a female’s lifetime, and we suppose that the number of female eggs is equal to the number of male eggs).

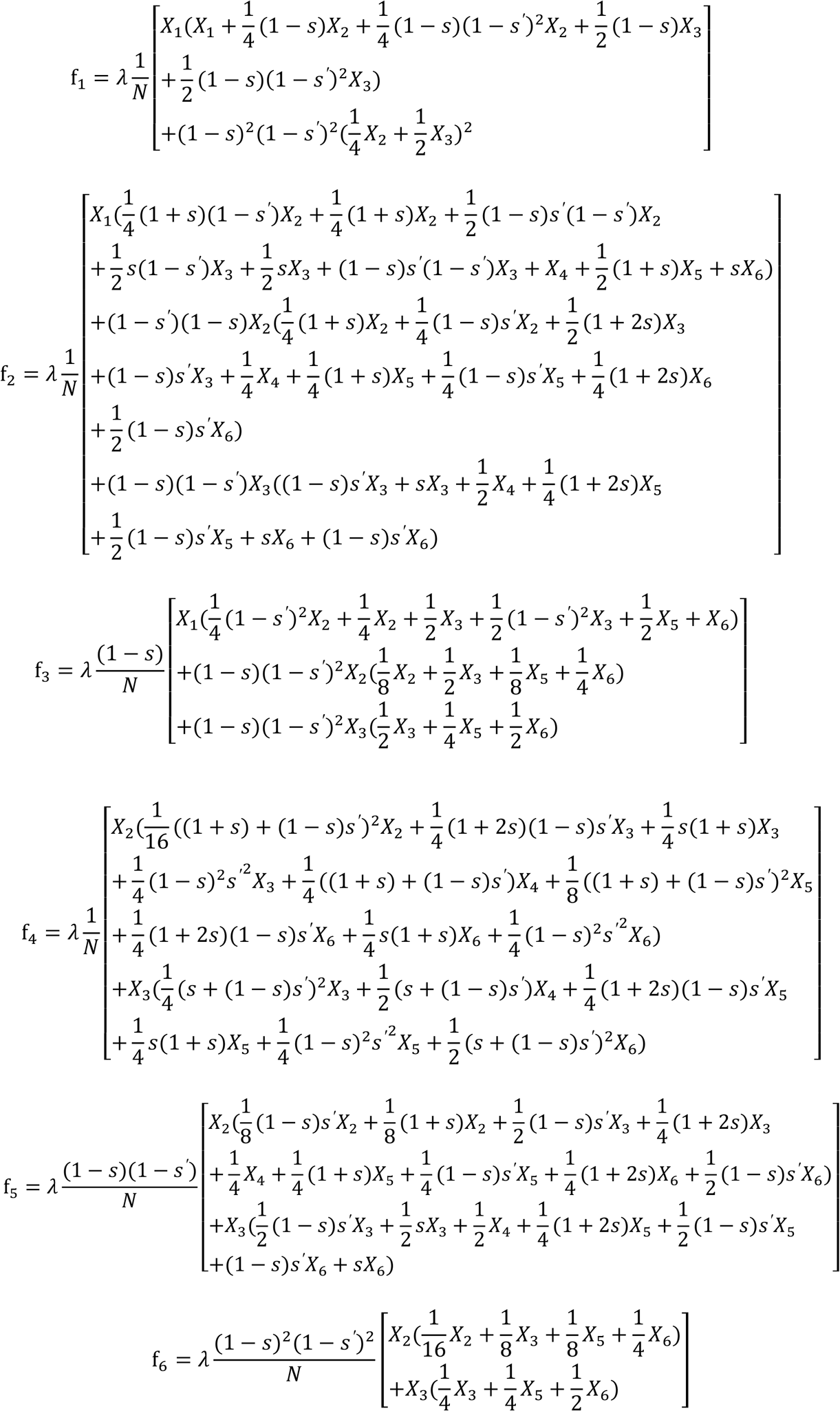

## References

1. Hay BA, Oberhofer G, Guo M. Engineering the composition and fate of wild populations with gene drive. Annu Rev Entomol, 66, 407–434, 2021.

2. Dhole S, Lloyd AL, Gould F. Gene drive dynamics in natural populations: The importance of density dependence, space, and sex. Annu Rev Ecol Evol Syst, 51, 505–531, 2020.

3. Rode NO, Estoup A, Bourguet D, Courtier-Orgogozo V, Débarre F. Population management using gene drive: molecular design, models of spread dynamics and assessment of ecological risks. Conserv. Genet. 20, 671–690, 2019.

4. Wang G-H, Du J, Chu CY, Madhav M, Hughes GL, Champer J. Symbionts and gene drive: two strategies to combat vector-borne disease. Trends Genet, 37, 708–723, 2022.

5. Hammond A, Galizi R, Kyrou K, Simoni A, Siniscalchi C, Katsanos D, Gribble M, Baker D, Marois E, Russell S, Burt A, Windbichler N, Crisanti A, Nolan T. A CRISPR-Cas9 gene drive system targeting female reproduction in the malaria mosquito vector Anopheles gambiae. Nat Biotechnol, 34, 78–83, 2016.

6. Kyrou K, Hammond AM, Galizi R, Kranjc N, Burt A, Beaghton AK, Nolan T, Crisanti A. A CRISPR-Cas9 gene drive targeting doublesex causes complete population suppression in caged Anopheles gambiae mosquitoes. Nat Biotechnol, 36, 1062–1066, 2018.

7. Carrami EM, Eckermann KN, Ahmed HMM, Sánchez C. HM, Dippel S, Marshall JM, Wimmer EA. Consequences of resistance evolution in a Cas9-based sex-conversion suppression gene drive for insect pest management. Proc Natl Acad Sci, 201713825, 2018.

8. Oberhofer G, Ivy T, Hay BA. Behavior of homing endonuclease gene drives targeting genes required for viability or female fertility with multiplexed guide RNAs. Proc Natl Acad Sci, 115, E9343–E9352, 2018.

9. Yang E, Metzloff M, Langmüller AM, Xu X, Clark AG, Messer PW, Champer J. A homing suppression gene drive with multiplexed gRNAs maintains high drive conversion efficiency and avoids functional resistance alleles. G3 Genes|Genomes|Genetics, 2022.

10. Champer SE, Oh SY, Liu C, Wen Z, Clark AG, Messer PW, Champer J. Computational and experimental performance of CRISPR homing gene drive strategies with multiplexed gRNAs. Sci Adv, 6, eaaz0525, 2020.

11. Hammond AM, Kyrou K, Bruttini M, North A, Galizi R, Karlsson X, Kranjc N, Carpi FM, D’Aurizio R, Crisanti A, Nolan T. The creation and selection of mutations resistant to a gene drive over multiple generations in the malaria mosquito. PLOS Genet, 13, e1007039, 2017.

12. Beaghton A, Beaghton PJ, Burt A. Gene drive through a landscape: Reaction-diffusion models of population suppression and elimination by a sex ratio distorter. Theor Popul Biol, 108, 51–69, 2016.

13. Beaghton A, Beaghton PJ, Burt A. Vector control with driving Y chromosomes: modelling the evolution of resistance. Malar J, 16, 286, 2017.

14. Simoni A, Hammond AM, Beaghton AK, Galizi R, Taxiarchi C, Kyrou K, Meacci D, Gribble M, Morselli G, Burt A, Nolan T, Crisanti A. A male-biased sex-distorter gene drive for the human malaria vector Anopheles gambiae. Nat Biotechnol, 38, 1054–1060, 2020.

15. Paril JF, Phillips BL. Slow and steady wins the race: Spatial and stochastic processes and the failure of suppression gene drives. Mol Ecol, 2022.

16. Birand A, Cassey P, Ross J V., Russell JC, Thomas P, Prowse TAA. Gene drives for vertebrate pest control: Realistic spatial modelling of eradication probabilities and times for island mouse populations. Mol Ecol, 31, 1907–1923, 2022.

17. Champer SE, Kim IK, Clark AG, Messer PW, Champer J. Anopheles homing suppression drive candidates exhibit unexpected performance differences in simulations with spatial structure. Elife, 11, 2022.

18. Liu Y, Champer J. Modelling homing suppression gene drive in haplodiploid organisms. Proc R Soc B, 289, 2022.

19. Champer J, Kim IK, Champer SE, Clark AG, Messer PW. Suppression gene drive in continuous space can result in unstable persistence of both drive and wild-type alleles. Mol Ecol, 30, 1086–1101, 2021.

20. Champer SE, Oakes N, Sharma R, García-Díaz P, Champer J, Messer PW. Modeling CRISPR gene drives for suppression of invasive rodents using a supervised machine learning framework. PLOS Comput Biol, 17, e1009660, 2021.

21. North AR, Burt A, Godfray HCJ. Modelling the suppression of a malaria vector using a CRISPR-Cas9 gene drive to reduce female fertility. BMC Biol, 18, 98, 2020.

22. North AR, Burt A, Godfray HCJ. Modelling the potential of genetic control of malaria mosquitoes at national scale. BMC Biol, 17, 26, 2019.

23. Beaghton AK, Hammond A, Nolan T, Crisanti A, Burt A. Gene drive for population genetic control: non-functional resistance and parental effects. Proceedings Biol Sci R Soc B Biol Sci, 286, 20191586, 2019.

24. Bull JJ, Remien CH, Krone SM. Gene-drive-mediated extinction is thwarted by population structure and evolution of sib mating. Evol Med public Heal, 2019, 66–81, 2019.

25. Liu Y, Teo W, Yang H, Champer J. Adversarial interspecies relationships facilitate population suppression by gene drive in spatially explicit models. bioRxiv, 2022.05.08.491087, 2022.

26. Piálek J, Barton NH. The spread of an advantageous allele across a barrier: The effects of random drift and selection against heterozygotes. Genetics, 145, 1997.

27. Barton NH, Hewitt GM. Analysis of hybrid zones. Annu Rev Ecol Syst, 16, 113–148, 1985.

28. Barton NH, Turelli M. Spatial waves of advance with bistable dynamics: Cytoplasmic and genetic analogues of allee effects. Am Nat, 178, E48–E75, 2011.

29. Tanaka H, Stone HA, Nelson DR. Spatial gene drives and pushed genetic waves. Proc Natl Acad Sci, 201705868, 2017.

30. Girardin L, Calvez V, Débarre F. Catch me if you can: A spatial model for a brake-driven gene drive reversal. Bull Math Biol, 2019.

31. Girardin L, Débarre F. Demographic feedbacks can hamper the spatial spread of a gene drive. J Math Biol, 83, 1–33, 2021.

32. Champer J, Lee E, Yang E, Liu C, Clark AG, Messer PW. A toxin-antidote CRISPR gene drive system for regional population modification. Nat Commun, 11, 1082, 2020.

33. Oberhofer G, Ivy T, Hay BA. Cleave and Rescue, a novel selfish genetic element and general strategy for gene drive. Proc Natl Acad Sci, 201816928, 2019.

34. Reeves RG, Bryk J, Altrock PM, Denton JA, Reed FA. First steps towards underdominant genetic transformation of insect populations. PLoS One, 9, e97557, 2014.

35. Maselko M, Feltman N, Upadhyay A, Hayward A, Das S, Myslicki N, Peterson AJ, O’Connor MB, Smanski MJ. Engineering multiple species-like genetic incompatibilities in insects. Nat Commun, 11, 1–7, 2020.

36. Buchman AB, Ivy T, Marshall JM, Akbari OS, Hay BA. Engineered reciprocal chromosome translocations drive high threshold, reversible population replacement in Drosophila. ACS Synth Biol, 7, 1359–1370, 2018.

37. Metzloff M, Yang E, Dhole S, Clark AG, Messer PW, Champer J. Experimental demonstration of tethered gene drive systems for confined population modification or suppression. BMC Biol, 20, 1–13, 2022.

38. Dhole S, Lloyd AL, Gould F. Tethered homing gene drives: A new design for spatially restricted population replacement and suppression. Evol Appl, eva.12827, 2019.

39. Weitzel AJ, Grunwald HA, Ceri W, Levina R, Gantz VM, Hedrick SM, Bier E, Cooper KL. Meiotic Cas9 expression mediates gene conversion in the male and female mouse germline. PLOS Biol, 19, e3001478, 2021.

40. Xu X, Harvey-Samuel T, Siddiqui HA, Ang JX De, Anderson ME, Reitmayer CM, Lovett E, Leftwich PT, You M, Alphey L. Toward a CRISPR-Cas9-based gene drive in the diamondback moth Plutella xylostella. Cris J, 5, 224–236, 2022.

41. Verkuijl SAN, Ang JXD, Alphey L, Bonsall MB, Anderson MAE. The challenges in developing efficient and robust synthetic homing endonuclease gene drives. Front Bioeng Biotechnol, 0, 426, 2022.

42. Champer J, Kim IK, Champer SE, Clark AG, Messer PW. Performance analysis of novel toxin-antidote CRISPR gene drive systems. BMC Biol, 18, 27, 2020.

43. Champer J, Champer SE, Kim IK, Clark AG, Messer PW. Design and analysis of CRISPR-based underdominance toxin-antidote gene drives. Evol Appl, eva.13180, 2020.

44. Zhu Y, Champer J. Simulations reveal high efficiency and confinement of a population suppression CRISPR toxin-antidote gene drive. bioRxiv, 2022.10.30.514459, 2022.

45. Champer J, Zhao J, Champer SE, Liu J, Messer PW. Population dynamics of underdominance gene drive systems in continuous space. ACS Synth Biol, 9, 779–792, 2020.

46. Langmüller AM, Champer J, Lapinska S, Xie L, Metzloff M, Champer SE, Liu J, Xu Y, Du J, Clark AG, Messer PW. Fitness effects of CRISPR endonucleases in Drosophila melanogaster populations. Elife, 11, 2022.

47. Champer J, Yang E, Lee E, Liu J, Clark AG, Messer PW. A CRISPR homing gene drive targeting a haplolethal gene removes resistance alleles and successfully spreads through a cage population. Proc Natl Acad Sci, 117, 24377–24383, 2020.

48. Pan M, Champer J. Making waves: Comparative analysis of gene drive spread characteristics in a continuous space model. bioRxiv, 2022.11.01.514650, 2022.

49. Li J, Champer J. Harnessing Wolbachia cytoplasmic incompatibility alleles for confined gene drive: a modeling study. bioRxiv, 2022.08.09.503337, 2022.

50. Kläy L, Girardin L, Calvez V, Débarre F. Pulled, pushed or failed: the demographic impact of a gene drive can change the nature of its spatial spread. arXiv, 2022.

51. Huang Y, Lloyd AL, Legros M, Gould F. Gene-drive into insect populations with age and spatial structure: a theoretical assessment. Evol Appl, 4, 415–428, 2011.

